# Signaling pathway screening platforms are an efficient approach to identify therapeutic targets in precision-medicine oriented early phase clinical trials

**DOI:** 10.1101/254342

**Authors:** Pedro Torres-Ayuso, Sudhakar Sahoo, Melanie Galvin, Hui Sun Leong, Kristopher K Frese, Andrew Hughes, Richard Marais, Caroline Dive, Matthew G Krebs, John Brognard

## Abstract

Precision medicine aims to tailor cancer therapies to target specific tumor-promoting aberrations. For tumors that lack actionable drivers, extensive molecular characterization and pre-clinical drug efficacy studies will be required to match patients with the appropriate targeted therapy. A cell line maintained at low passage and a patient-derived xenograft model (PDX) were generated using a fresh biopsy from a patient with a poorly-differentiated neuroendocrine tumor of unknown primary origin. Next-generation sequencing, high throughput signaling network analysis, and drug efficacy trials were then conducted to identify actionable targets for therapeutic intervention. No actionable mutations were identified after whole exome sequencing of the patient’s DNA; however, whole genome sequencing revealed amplification of the 3q and 5p chromosomal arms, that include the *PIK3CA* and *RICTOR* genes, respectively. Consistent with amplification of these genes, pathway analysis revealed activation of the AKT pathway. Based on this analysis, efficacy of PIK3CA and AKT inhibitors were evaluated in the tumor biopsy-derived cell culture and PDX, and response to the AKT inhibitor AZD5363 was observed both *in vitro* and *in vivo* indicating the patient would benefit from targeted therapies directed against the serine/threonine kinase AKT. In conclusion, our study demonstrates that high throughput signaling pathway analysis complements next-generation sequencing approaches for detection of actionable alterations and will aid in patient stratification into early-phase clinical trials.

## Introduction

The application of precision medicine into clinical practice has revolutionized the management of cancer over the last decade. Next-generation sequencing (NGS) technologies allow the identification of actionable mutations in tumors, to which targeted therapies can be developed with the potential to improve therapeutic index regarding impact on tumor versus normal tissues in contrast to conventional cytotoxics. Increasingly, NGS of patient tumor samples guides patient stratification into clinical trials, such that only the patients bearing specific molecular alterations will receive the corresponding targeted therapy.

The TARGET (**<u>T</u>**umour ch**<u>AR</u>**acterisation to **<u>G</u>**uide **<u>E</u>**xperimental Targeted **<u>T</u>**herapy) protocol aims to stratify patients based on genetic alterations identified in tumor specimens and/or circulating-free DNA. In addition, *in vivo* drug efficacy studies in patient-derived xenografts (PDX) are performed for a subset of patients. This approach matches patients to the most appropriate and available early phase clinical trials according to their molecular alterations to maximize the chance of patient benefit from targeted therapies^1^.

Here we describe a case study (TAR007) of a patient with no smoking history and a poorly differentiated neuroendocrine tumor of unknown primary origin. These are rare tumors, characterized by poor prognosis, and these patients have limited treatment options. Chemotherapy and/or radiotherapy treatment, prior to or after surgical resection of detected tumor masses may be utilized, but there is limited data for the use of targeted therapies to treat this type of cancer^2^. In this study, we conducted an extensive molecular characterization of a freshly resected tumor biopsy to identify constitutively activated and druggable cell survival pathways in this tumor specimen. While whole exome sequencing (WES) did not show any actionable mutations, *PIK3CA* and *RICTOR* gene amplifications were detected by whole genome sequencing (WGS). Complementing this approach, we used a high-throughput platform for analysis of cell signaling pathways and detected hyperactivation of the AKT signaling axis. Treatment of tumor biopsy-derived cell cultures, or a successfully established PDX model showed response to AKT inhibitors, and little or no effect of PI3K inhibitors. These results highlight that combining NGS, signaling pathway analyses, and preclinical drug efficacy studies can successfully identify activated pathways that can be targeted therapeutically. In addition, we identify amplified *PIK3CA* and *RICTOR* as potential biomarkers for patients with neuroendocrine tumors with increased propensity to respond to treatment with AKT inhibitors.

## Results

A patient presented with a poorly differentiated neuroendocrine tumor of unknown primary origin, and the tumor was surgically resected from the right axilla. An experimental pre-clinical study was designed to identify potentially druggable genetic alterations (Figure 1A). Tumor samples were inoculated into NSG mice to generate a PDX mouse model for use in drug efficacy studies. In parallel, tumor fragments were frozen for DNA sequencing and a low passage cell culture was derived after digestion of the tumor specimen.

**Figure 1.**
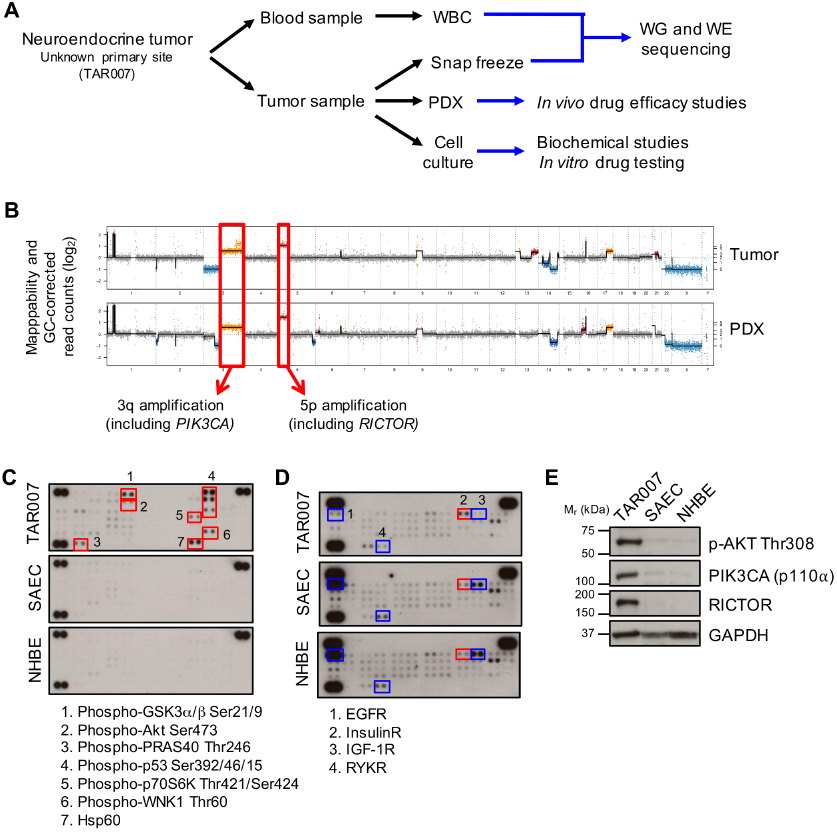
Activation of the AKT pathway in a neuroendocrine tumor of unknown primary origin. (A) Scheme depicting sample collection and processing for the TARGET clinical trial and the studies performed for the molecular analysis of the tumor specimen TAR007. WBC: White Blood Cells; WG: Whole Genome; WE: Whole Exome. (B) Whole genome sequencing from the tumor and a patient-derived sample shows extensive copy number variations, including amplifications in 3q and 5p chromosomes. (C) Phosphokinase arrays from tumor-derived low passage cell cultures show activation of several components of the AKT-mTOR signaling pathway compared to normal primary cells NHBE (normal human bronchial epithelium) and SAEC (small airway epithelial cells). (D) Phospho-receptor tyrosine kinase (RTK) arrays show different RTK activation profiles. (E) Western blot confirming overexpression of PIK3CA (aka p110α) and RICTOR, as well as hyperactivation of AKT in TAR007-derived cell cultures. GAPDH was used as a loading control.

WES of the tumor specimen was conducted to identify targetable alterations in this patient’s tumor, and DNA from white blood cells was used as a control to exclude germline variants from the analysis. WES did not reveal any actionable mutations (Supplemental file 1). WGS was then performed to identify copy number alterations. WGS analysis from both the tumor specimen and a PDX-derived sample revealed chromosome deletions at 3p and 14q, and amplifications in 3q, 5p, 13q and 17q (Figure 1B). Among the regions of amplification, we detected copy number gains of the *PIK3CA* (in 3q, 4 copies) and *RICTOR* (in 5p, 5 copies) genes, which are upstream activators of the oncogenic kinase AKT.

Fresh tumor biopsy-derived low passage cell cultures were then used to assess pathways activated in the tumor sample. Normal, non-transformed, primary cells were used as controls (NHBE and SAEC; Figure 1C). Several components of the AKT pathway were specifically activated in the tumor-derived cells, including AKT, PRAS40, and GSK3α/β, consistent with this pathway being activated due to amplification of *RICTOR* and *PIK3CA*. High levels of phosphorylation were also found for the p53 tumor suppressor protein, and the WNK1 protein kinase (Figure 1C). Using phospho-receptor tyrosine kinase (RTK) arrays, hyperactivation of the insulin receptor was detected. Interestingly, we observed downregulation of the epithelial growth factor (EGF), insulin-like growth factor 1 (IGF-1) and Receptor-Like Tyrosine Kinase (RYK) receptors (Figure 1D). Overexpression of PIK3CA, RICTOR, and activation of the AKT pathway, were further confirmed by western blot (Figure 1E).

Our data indicated that the PI3K/AKT pathway was constitutively activated so inhibitors targeting these kinases were explored in functional assays. Treatment of the tumor-derived cells with AZD8835 or GDC0941, two PIK3CA selective inhibitors, had minor pro-apoptotic effects compared to vehicle-treated cells (Figure 2A and S1A). In contrast, treatment of the tumor-derived cells with two selective AKT inhibitors, AZD5363 (an ATP-competitive inhibitor), or MK2206 (an allosteric AKT inhibitor) resulted in a significant increase in cell death (Figure 2A and S1A). In addition, AKT inhibition had a strong effect on cell proliferation, and diminished the percent of proliferating cells (21.1% in vehicle-treated cells; 16.0% in AZD8835-treated cells and 9.0% in AZD5363-treated cells; Figure 2B). Time course experiments were then conducted to investigate mechanisms of sensitivity to AKT inhibitors. Treatment with the AKT inhibitors, AZD5363 or MK2206, resulted in sustained inactivation of the AKT substrate PRAS40, this was not the case with PI3K inhibitors (BKM120, GDC0941 and AZD8835; Figure 2C, D and S1B). PRAS40 is a negative regulator of mTORC1, a multiprotein complex required for cell growth and proliferation. Phosphorylation of PRAS40 by both AKT and mTORC1 promotes the dissociation of PRAS40 from mTOR, leading to activation of mTORC1 (Figure 2C)^3^. Decreased phosphorylation of the mTORC1 downstream substrate ribosomal protein S6 (rpS6) was also observed in cells treated with AKT inhibitors, consistent with nonphosphorylated PRAS40 binding and inhibiting mTORC1 activation (Figure S1B). Cells treated with PI3K inhibitors did not display sustained decreased rpS6 phosphorylation (Figure S1B). Therefore, sustained inactivation of pathways downstream of AKT in cells treated with AKT inhibitors is likely to account for the sensitivity observed towards the AKT inhibitors and not to PI3K inhibitors.

**Figure 2.**
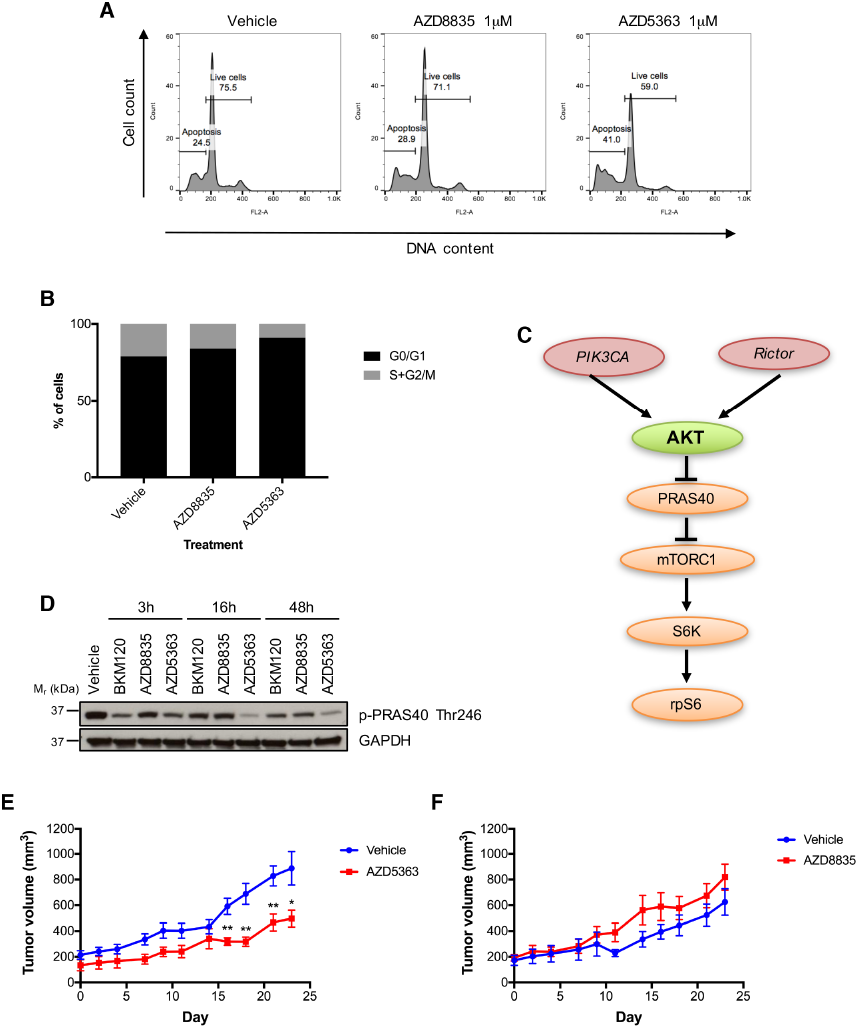
TAR007 patient-derived cancer cells and xenografts (PDX) are sensitive to the AKT inhibitor AZD5363. (A) Tumor-derived low passage cells were treated with the indicated inhibitors for 48h and DNA content was analyzed by propidium iodide staining and flow cytometry. The percentages of live and dead cells are indicated. *N* = 3 experiments. (B) Percentage of cells in each phase of the cell cycle (G0/G1 or S+G2/M) of cells analyzed as in A. (C) Scheme of the AKT pathway. (D) TAR007 cells were treated at 1μM concentration with the indicated inhibitors for different times and effectors of the AKT-mTOR pathway were analyzed by western blot. GAPDH was used as a loading control; *N* = 3 experiments. (E, F) TAR007 PDX (*N*=9 mice/group) were treated with (E) vehicle and AZD5363 or (F) vehicle and AZD8835 as described in the methods section, for the days indicated and tumor volume was recorded every 48h. Data are shown as mean ± SEM. ** *P*-value<0.01; * *P-*value<0.05.

We then expanded these *in vitro* studies to explore responses in PDX models. Consistent with data derived from the low passage cell cultures, treatment with AZD5363 *in vivo* significantly reduced tumor growth by approximately 50% (Figure 2E). PI3K inhibition with AZD8835 showed no significant difference in the tumor growth rate when compared with the vehicle-treated group (Figure 2F). Together our data confirm inhibition of AKT as a viable therapeutic strategy and demonstrate WES, WGS, molecular pathway analysis, and PDX mouse models can be used in combination to significantly aid in stratifying patients for treatment with targeted therapies.

## Discussion

Here, we report extensive molecular characterization from a high grade neuroendocrine tumor of unknown primary origin with the aim of identifying actionable alterations. Increasingly, NGS approaches, such as WES, are being used in the clinical setting to facilitate identification of actionable mutations. However, in this patient case study, WES analysis of the surgical specimen did not reveal the presence of any targetable genetic alterations. Amplification of chromosomes 3q and 5p were identified by WGS; this included copy number gains in the *RICTOR* and *PIK3CA* genes.

High throughput technologies analyzing cellular signaling networks, including phospho-arrays, are being implemented into therapeutic target and biomarker discovery programs^4,5^. These approaches aid in identifying signaling networks activated in tumor specimens, as genetic mutations do not always correlate with changes in protein activity. Phospho-array analysis revealed hyperactivation of the PI3K/AKT pathway, consistent with amplification of the *RICTOR* and *PIK3CA* genes. Both PI3K and RICTOR are upstream activators of AKT and were confirmed to be overexpressed at the protein level.

A major strength of the TARGET protocol is the ability to simultaneously generate patient-derived preclinical models in which to test the efficacy of potential personalized treatments, so that a patient may be enrolled onto the trial most likely to offer clinical proof of concept and patient benefit. PDX mouse models are to date considered to accurately predict tumor responses, however, it can take a prolonged time to expand the tumor specimen in animals before any pharmacological study can be conducted. To overcome this issue, we also established low passage tumor derived cell cultures to test the efficacy of PI3K and AKT inhibitors based on the molecular alterations detected. Increased cell death was observed after treatment with the AKT inhibitors AZD5363 and MK2206, however, little or no response was observed to PI3K inhibitors. This is in agreement with recent data, showing that tumors with *RICTOR* amplification are frequently refractory to PI3K inhibition^6^. Sensitivity to AKT inhibitors might rely on sustained inhibition downstream of AKT and mTORC1, an effect not observed after PI3K inhibition. Patients with *RICTOR* amplification might also benefit from mTOR inhibitors^6^; however, these compounds were not tested since secondary resistance normally appears and involves reactivation of the PI3K-AKT or the MAPK pathways^7,8^. Similar responses to AKT and PI3K inhibitors were observed in our PDX models, indicating that low passage cell cultures, when possible to generate, can be a powerful tool for initial inhibitor testing and aid in refining *in vivo* experiments. In addition, our data suggest that *PIK3CA* and *RICTOR* gene co-amplification may define a unique subset of patients with neuroendocrine tumors that will respond to AKT inhibitors, and could thus be used as an additional biomarker for treatment stratification.

In summary, we show development of a precision medicine approach to aid in stratification of patients into early phase clinical trials. To date enrollment into clinical trials mostly relies on mutational status of the corresponding target (e.g. AKT). Our study suggests that complementation of sequencing approaches with cost-effective and time-efficient antibody-based technologies, such as phospho-arrays, can maximize the detection of druggable pathways in tumor cases where no actionable mutations have been identified or little is known about tumor etiology. Clinical implementation of these unbiased signaling pathway analysis technologies could improve selection of patients to early phase clinical trials of targeted therapies, and benefit a patient cohort that lacks targetable genetic alterations. Integration of these technologies into the clinic will aid in precision medicine based approaches.

## Methods

### Patient details

A 68-year old gentleman presented with an isolated right axillary mass in October 2014. An axillary biopsy revealed poorly differentiated (Grade 3, Ki67 85%) neuroendocrine carcinoma of unknown origin (positive for CD56, chromagranin and synaptophysin; negative for TTF1, CDX2 and Merkel Cell Polyomavirus). Positron emission technology/computed tomography demonstrated a right axillary mass and no other identifiable sites of disease. The patient received 6 cycles of carboplatin and etoposide chemotherapy between Oct 2014 and April 2015 with RECIST partial response after 3 cycles but with evidence of tumour growth after cycle 6. He was referred to the Experimental Cancer Medicine Team and consented to TARGET in June 2015. In parallel he was referred for surgical resection of the isolated axillary mass which achieved a complete resection and permitted access to fresh tissue for PDX and translational research in July 2015.

### Phospho-arrays

The Proteome Profiler Human Phospho-Kinase and Human Phospho-Receptor Tyrosine Kinase (RTK) Array kits (R&D Systems) were used according to the manufacturer’s instructions.

### Statistical analysis

Statistical analysis was done with GraphPad Prism, the Mann-Whitney test was used to assess differences between treatments; when *P*-value < 0.05, differences were considered significant.

Additional methods are described in the supplemental material.

## Data availability statement

All data generated or analyzed during this study are included in this published article and its supplementary information files.

## Acknowledgements

We thank the patient and his family for donating samples for this research. AZD5363 and AZD8835 were kindly donated by AstraZeneca. We thank members of the Signalling Networks in Cancer team, the Clinical and Experimental Pharmacology *in vivo* team, Dr MR Girotti and Dr F Trapani at the Cancer Research UK Manchester Institute for helpful discussions. We thank the Biological Resources Unit, Molecular Biology and Flow Cytometry Core Facilities at Cancer Research UK Manchester Institute for technical assistance, and the Manchester Cancer Research Centre Biobank for logistical support. We also thank the National Institute for Health (NIHR) Manchester Clinical Research Facility.

## Contributions

PT-A, MGK and JB conceived, designed and supervised the study; PT-A performed and analyzed the experiments; SS and HSL Leong analyzed the sequencing data; KKF, MG and CD assisted in the *in vivo* studies; PT-A and JB interpreted the data, wrote and prepared the manuscript; KKF, CD and MGK revised the manuscript. AH, RM, CD and MGK conceived and set the TARGET trial protocol.

## References

1 Krebs, M. et al. TARGET trial: Molecular profiling of circulating tumour DNA to stratify patients to early phase clinical trials. Journal of Clinical Oncology 34, TPS11614–TPS11614, doi:10.1200/JCO.2016.34.15_suppl.TPS11614 (2016).

2 Spigel, D. R., Hainsworth, J. D. & Greco, F. A. Neuroendocrine carcinoma of unknown primary site. Seminars in oncology 36, 52-59, doi:10.1053/j.seminoncol.2008.10.003 (2009).

3 Laplante, M. & Sabatini, D. M. mTOR signaling in growth control and disease. Cell 149, 274–293, doi:10.1016/j.cell.2012.03.017 (2012).

4 Masuda, M. & Yamada, T. Signaling pathway profiling by reverse-phase protein array for personalized cancer medicine. Biochim Biophys Acta 1854, 651–657, doi:10.1016/j.bbapap.2014.10.014 (2015).

5 Lu, Y. et al. Using reverse-phase protein arrays as pharmacodynamic assays for functional proteomics, biomarker discovery, and drug development in cancer. Seminars in oncology 43, 476–483, doi:10.1053/j.seminoncol.2016.06.005 (2016).

6 Cheng, H. et al. RICTOR Amplification Defines a Novel Subset of Patients with Lung Cancer Who May Benefit from Treatment with mTORC1/2 Inhibitors. Cancer Discov 5, 1262–1270, doi:10.1158/2159-8290.CD-14-0971 (2015).

7 O’Reilly, K. E. et al. mTOR inhibition induces upstream receptor tyrosine kinase signaling and activates Akt. Cancer Res 66, 1500–1508, doi:10.1158/0008-5472.CAN-05-2925 (2006).

8 Carracedo, A. et al. Inhibition of mTORC1 leads to MAPK pathway activation through a PI3K-dependent feedback loop in human cancer. J Clin Invest 118, 3065–3074, doi:10.1172/JCI34739 (2008).

